# Predicting vertical and shear ground reaction forces during walking and jogging using wearable plantar pressure insoles

**DOI:** 10.1101/2023.02.19.529141

**Authors:** Maryam Hajizadeh, Allison L. Clouthier, Marshall Kendall, Ryan B. Graham

## Abstract

**Background:** The development of plantar pressure insoles has made them a potential replacement for force plates. These wearable devices can measure multiple steps and might be used outside of the lab environment for rehabilitation and evaluation of sport performance. However, they can only measure the normal force which does not completely represent the vertical ground reaction force (GRF). In addition, they are not able to measure shear forces which play an import role in the dynamic performance of individuals. Indirect approaches might be implemented to improve the accuracy of the force estimated by plantar pressure systems.

**Research question:** The aim of this study was to predict the vertical and shear components of ground reaction force from plantar pressure data using recurrent neural networks.

**Methods:** GRF and plantar pressure data were collected from sixteen healthy individuals during 10 trials of walking and five trials of jogging using Bertec force plates and FScan plantar pressure insoles. A long short-term memory (LSTM) neural network was built to consider the time dependency of pressure and force data in predictions. The data were split into three subsets of train, to train the LSTM model, evaluate, to optimize the model hyperparameters, and test sets, to assess the accuracy of the model predictions.

**Results:** The results of this study showed that our LSTM model could accurately predict the shear and vertical GRF components during walking and jogging. The predictions were more accurate during walking compared to jogging. In addition, the predictions of mediolateral force had higher error and lower correlation compared to vertical and anteroposterior components.

**Significance:** The LSTM model developed in this study may be an acceptable option for accurate estimation of GRF during outdoor activities which can have significant impacts in rehabilitation, sport performance, and gaming.

## 1. Introduction

Recent developments in plantar pressure insoles have increased their potential to provide real-time information about forces applied to the foot during outdoor activities. An array of resistive, capacitive, or piezoelectric sensors are integrated within these insoles [1-3] to measure the pressure perpendicular to the plane of the sensors and subsequently the total force. The advantage of these wearable devices over traditional instruments such as force plates is that they are wearable and can monitor multiple steps, while imposing no constraint on foot placement. In addition, they can provide real-time feedback on peoples’ everyday health or athletic performance and can be used to evaluate and optimize the performance of footwear and orthotics [4]. However, the total force computed from these devices is less accurate than the gold standard vertical ground reaction force (GRF) recorded from force plates [5]. Several factors might contribute to lower accuracy of these systems. Primarily, the total force calculated from plantar pressure insoles is a combination of vertical and shear forces applied to the subsurface of the foot due to the distortion of insoles with the curvatures of the foot and shoe sole [6-8]. However, the complexities of the soft tissues in the foot surface make it hard to extract shear forces from plantar pressure data [6, 8]. As a result, the total force is presented as vertical GRF. Currently, anteroposterior (AP) and mediolateral (ML) GRFs, which are crucial components for diagnosis and treatment of pathological symptoms and sport injuries, cannot be measured by pressure insoles. Other factors contribute to inaccuracy and variability including the motion artifact of insoles during dynamic activities, calibration problems, and sensor limitations such as drift and hysteresis [5, 9-11].

Indirect approaches have been suggested to enhance the accuracy of total force estimations and predict the 3D components of GRF from commercial pressure insoles. Previous research has recommended to apply further calibrations in pressure measurements to reduce the effect of motion artifact and sensor limitations, but this was only partially successful [12]. Alternatively, statistical tools and neural networks have been used to estimate accurate GRFs. Fong *et al*. [13] applied stepwise linear regression analysis to extract the three components of GRF from Novel’s Pedar pressure insoles, containing an array of 99 capacitive sensors, during walking. The estimated force showed very good correlation to force plate measures for AP (0.928) and vertical (0.989) components, but lower correlation for ML force (0.719) [13]. Another study that proposed multi-stage regression to calculate forces from Pedar pressure insoles during walking reported the root mean square error normalized to peak recorded values (NRMSE) as of 21% for ML force, 10% for AP force, and 3% for vertical forces [14]. Rouhani *et al*. [15] implemented principal component analysis to remove the redundancy of pressure data from Pedar insole data followed by a locally linear neuro-fuzzy model for predicting GRF. Their model could consider the non-linear mapping between pressure and GRF in contrast to regression models. The results of their study showed the NRMSE as of 21.58 %, 17.32%, and 15.51% for ML, AP, and vertical GRF components, respectively [15]. Savelberg *et al*. [16] trained an artificial neural network using a supervised learning method with backpropagation to predict the AP force from Novel’s Emed plantar pressure data. They reported different levels of model performances, ranging from poor to good, for inter-subject predictions, which limits the robustness and practicality of their method [16]. A further study found better accuracy and correlation for GRF prediction with the same plantar pressure insoles using principal component analysis-mutual information and wavelet neural networks, which can consider the local data structure and non-linear dependencies. The NRMSE reported in that study was 15.027% for ML, 12.920% for AP, and 12.957% for vertical forces [17]. Mathematical techniques have been applied as a further solution to calculate accurate GRF from pressure data. DeBerardinis *et al*. [5] used transfer functions to improve the accuracy of vertical GRF measurements from Medilogic plantar pressure insoles for each insole size. Their model could reduce the RMSE for vertical GRF to 10% of body weight, but they reported that transfer functions did not consistently improve the GRF calculations. Therefore, more robust techniques with higher levels of accuracy are demanded in this domain to reach the maximum potential of plantar pressure insoles.

The focus of this work was to assess the ability of recurrent neural networks to accurately predict vertical and shear components of GRF from Tekscan’s F-Scan system. F-Scan systems, which constitute a collection of resistive sensels, have cost advantages over Pedar systems with capacitive sensels. However, the accuracy of systems with resistive sensels has been stated to be significantly affected by hysteresis, drift and variability [12]. Therefore, developing predictive approaches that are capable to estimate accurate GRF from Tekscan plantar pressure systems would make these devices more practical in research, rehabilitation, and sport applications. To our knowledge, previous research using neural networks to predict GRF from plantar pressure systems did not fully consider the time-dependency and sequencing of pressure and GRF during gait for predicting vertical and shear components of force. Therefore, the first objective of this study was to determine the validity of F-Scan insoles for predicting 3D GRFs using recurrent neural networks during walking. The second objective of this study was to compare the accuracy of predicted GRFs between walking and jogging as higher shear and bending loads are applied to the sensors in jogging. Long short-term memory (LSTM) networks were used in this study to consider the time-dependency and patterns of pressure and force data.

## 2. Methods

### 2.1. Experimental Procedure

Sixteen participants (7 females and 9 males, 25.1±2.9 years, 1.73±0.99 m, 72.9±14.1 kg) were recruited for two testing sessions (1 week apart) and provided informed consent for this research which was approved by the University of Ottawa. Plantar pressure insoles (3000E, F-Scan, Tekscan, USA) were cut to fit into the participant’s comfortable shoes. In each session, the participants were asked to do 10 walking and five jogging trials at a self-selected pace where their dominant foot had complete contact with a force plate (FP-4060, Bertec, USA) in the middle of the walkway. Motion capture software (Nexus 2.5, Vicon, UK) was used to collect force plate data (1000 Hz), and Tekscan software recorded the pressure data simultaneously at 100 Hz. A trigger was used to synchronize the force plate and pressure data.

### 2.2. Data Processing

The primary data preparation was performed in MATLAB (2018b, MathWorks, USA) using Tekscan’s software developer toolkit to read pressure files and ezc3d to read force plate data [18]. The 955 individual sensels in F-Scan were organized into a 60*21 pressure matrix and sensel dropouts were addressed by replacing the missed sensel with average of its neighboring sensels. Then, the size of the pressure matrix was reduced by averaging the pressure value of each group of 4*4 adjacent sensels to create the masked pressure data (Supplementary Figure 1). These masked pressure data included 252 pressure data points and were used for further analysis. Both force plate and pressure trials were then amplitude normalized to the participant’s body weight and maximum pressure, respectively, and time normalized to 101 points representing the entire stance phase. These trials were finally concatenated for all participants to create matrices that the neural network would use as input and targets. Due to bad data during collections, two walking trials and three jogging trials had to be removed resulting in a matrix of 32118× 252 for walking and matrix of 15352× 252 for jogging. A detailed description of these matrix structures is presented in Table 1.

**Table 1.** Structure of LSTM model.

|  | Walking | Jogging |
| --- | --- | --- |
| Input feature | Pressure values of masked pressure matrix (# 252) | Pressure values of masked pressure matrix (# 252) |
| Data division | In total: 32 subjects, 10 trials of each subject, 100% normalized to stance phase ($32 \times 10 \times 101 - 2 \times 101 = 32118$ data points)<br>- Training (22 subjects=22119 datasets)<br>- Validation (5 subjects=5050 datasets)<br>- Test (5 subjects=4949 datasets) | In total: 31 subjects, 5 trials of each subject, 100% normalized to stance phase ($31 \times 5 \times 101 - 3 \times 101 = 15352$ data points)<br>- Training (21 subjects=10403 datasets)<br>- Validation (5 subjects=2525 datasets)<br>- Test (5 subjects=2424 datasets) |
| Output feature | GRF-x (#1)<br>GRF-y (#1)<br>GRF-z (#1) | GRF-x (#1)<br>GRF-y (#1)<br>GRF-z (#1) |
| Neural net structure (with tuned hyperparameters) | *GRF-x:*<br>- 1 input layer (252 cells)<br>- 4 LSTM layers (64 cells, dropout=0.247)<br>- 1 fully connected layer (1 cell)<br>- Learning rate=0.006<br>*GRF-y:*<br>- 1 input layer (252 cells)<br>- 2 LSTM layers (64 cells)<br>- 1 fully connected layer (1 cell)<br>- Learning rate=0.003<br>*GRF-z:*<br>- 1 input layer (252 cells)<br>- 3 LSTM layers (128 cells)<br>- 1 fully connected layer (1 cell)<br>- Learning rate=0.01 | *GRF-x:*<br>- 1 input layer (252 cells)<br>- 4 LSTM layers (256 cells, dropout=0.765)<br>- 1 fully connected layer (1 cell)<br>- Learning rate=4.2e-5<br>*GRF-y:*<br>- 1 input layer (252 cells)<br>- 3 LSTM layers (128 cells)<br>- 1 fully connected layer (1 cell)<br>- Learning rate=0.01<br>*GRF-z:*<br>- 1 input layer (252 cells)<br>- 2 LSTM layers (256 cells, dropout=0.147)<br>- 1 fully connected layer (1 cell)<br>- Learning rate=0.01 |

### 2.3. Artificial neural network model

Since the prediction of GRF using plantar pressure data was a sequential problem and considering temporal information was important, LSTM networks were used for predictions. Progressive windows of pressure data were used to predict each GRF component at each time frame of stance phase, as shown in Figure 1. A time step period of pressure data (101 data points, red shadow) was taken as input to predict a single data point of GRF (red spot) as target. Then, the window for pressure data was shifted forward frame by frame (1 data point) to finish the prediction of GRF during the whole stance phase. The time step period for pressure data was regarded as 101 data points equal to a whole period of stance phase. The data were divided in training, validation, and tests sets as shown in Table 1. The neural networks were implemented in PyTorch and were composed of LSTM with hidden layers and dropout layer to avoid overfitting, followed by a fully connected dense layer to output predictions. An adaptive moment estimation (Adam) optimizer was used to train the neural network with a mean squared error (MSE) loss criterion. The number of LSTM layers, number of LSTM cells, LSTM dropout percentage, and the learning rate of the optimizer were considered as hyperparameters and tuned through Bayesian hyperparameter optimization using the Ax Platform [19, 20]. The mean square error from the predictions of the validation data set was used as the criterion for optimization. Through this analysis, optimal parameters for each component of GRF during both walking and jogging were found as reported in Table 1. In addition, the number of epochs and batch size were set to 25 and 101, respectively, for all the models.

**Figure 1.**
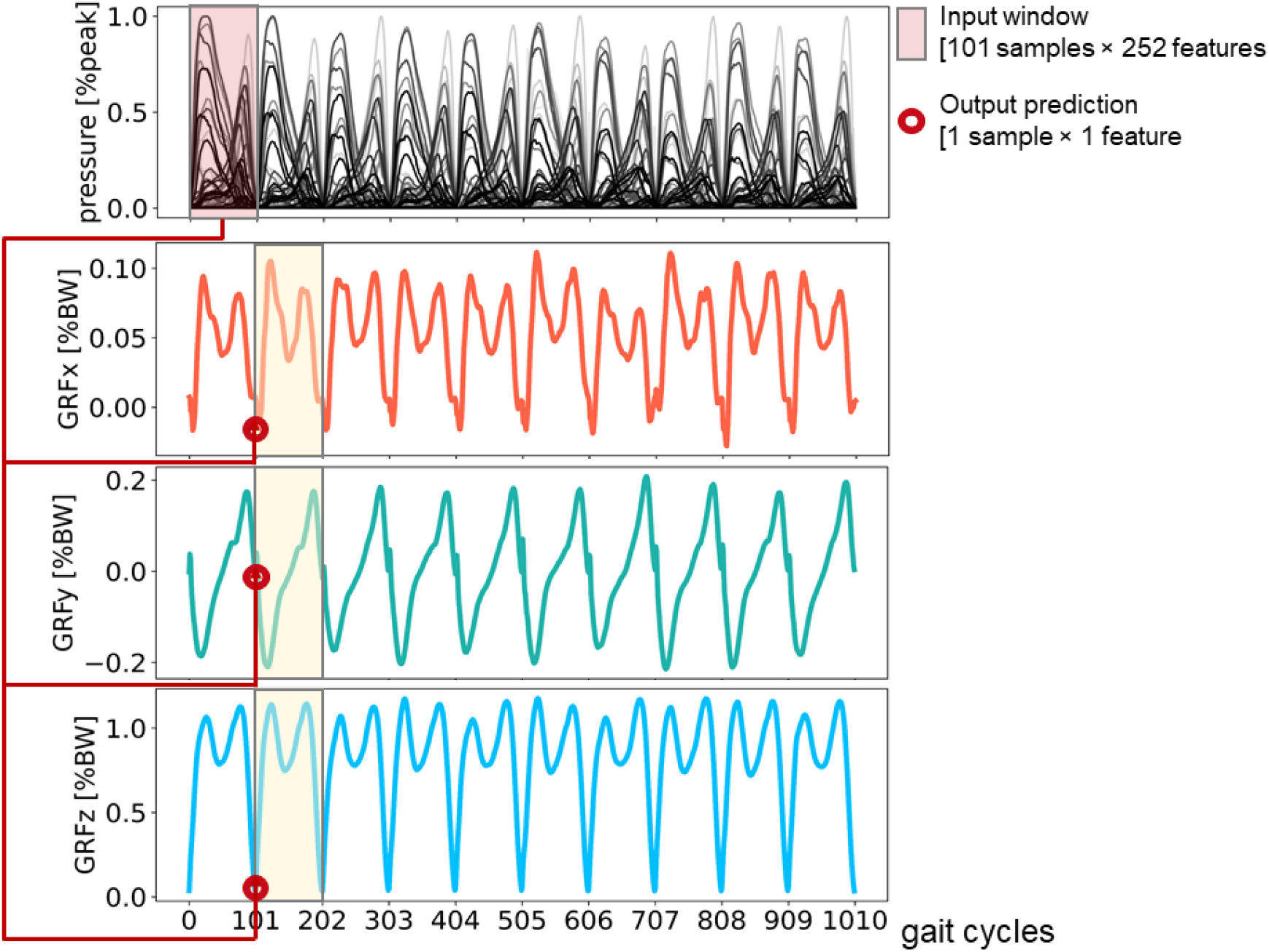
An example illustrating the prediction of three components of ground reaction force from plantar pressure data using long short-term memory model. The pink shadow is the input feature data. The red spots represent the corresponding predicted spots of force. The yellow shadow is the ground truth for ground reaction force. GRF-ML represents the mediolateral force, GRF-AP represents the anteroposterior force, and GRF-vertical represents the vertical force

### 2.4. Statistical analysis

To evaluate the performance of the LSTM model, four parameters were computed to analyze the accuracy of predictions for the trajectories of GRFs (ML, AP, vertical) across the test data set: (1) mean absolute error (MAE), (2) Root Mean Square Error (RMSE), (3) normalized root mean square error (NRMSE) defined as RMSE normalized to the range of GRF component, and (4) Pearson correlation coefficient. The mean and standard deviation of error and correlation coefficients were estimated across the different trials of each participant in the test set as well as across all the available trials in the test set for walking and jogging activities.

## 3. Results

Tuning the hyperparameters resulted in different model parameters for predicting GRF components during walking and jogging. These hyperparameters were summarized in Table 1 and were used to predict the trajectories of three GRF components from pressure data in test dataset. Figure 2 shows the mean predicted trajectories across all participants in the test set compared to GRF components obtained from force plates for both walking and jogging sessions. In addition, the predicted GRF components were compared to the measured data for each participant in test set in supplementary Figures 2-4 for walking and in supplementary Figures 5-7 for jogging. The pattern of predicted forces was more consistent with force plate data for participants 2 and 5 compared to other participants during walking. However, similar levels of consistency were observed between all participants during jogging.

**Figure 2.**
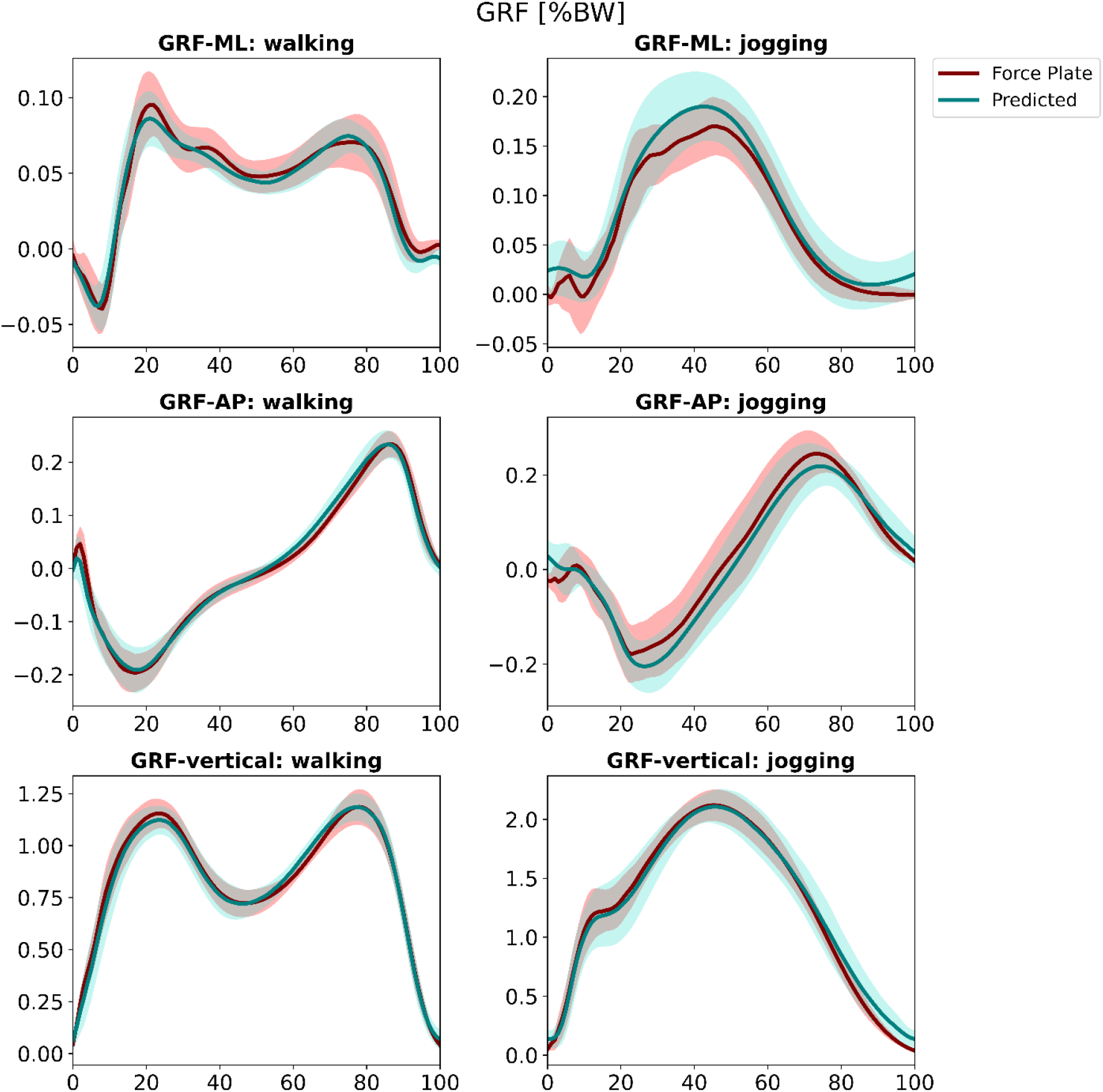
The ground truth and predicted mediolateral, anteroposterior, and vertical components of ground reaction force during walking and jogging. The forces are reported as a percentage of body weight.

The model performance for predicting the GRF components was estimated and shown in Figures 3-4. Figure 3 shows MAE, RMSE, and NRMSE and Figure 4 shows the correlation coefficients between the predicted force and the measured force in the test set for each participant as well as for all participants combined during walking and jogging. The average errors during walking were MAE= 5.22± 2.05% of body weight and NRMSE= 5.50± 2.25% for vertical GRF, MAE = 2.10± 0.86% of body weight and NRMSE= 6.26± 2.52% for AP GRF, and MAE= 1.48± 0.45% of body weight and NRMSE= 12.96± 4.98% for ML force. The average correlation coefficient was 0.985± 0.010 for vertical force, 0.983± 0.018 for AP force, and 0.926± 0.062 for ML force.

**Figure 3.**
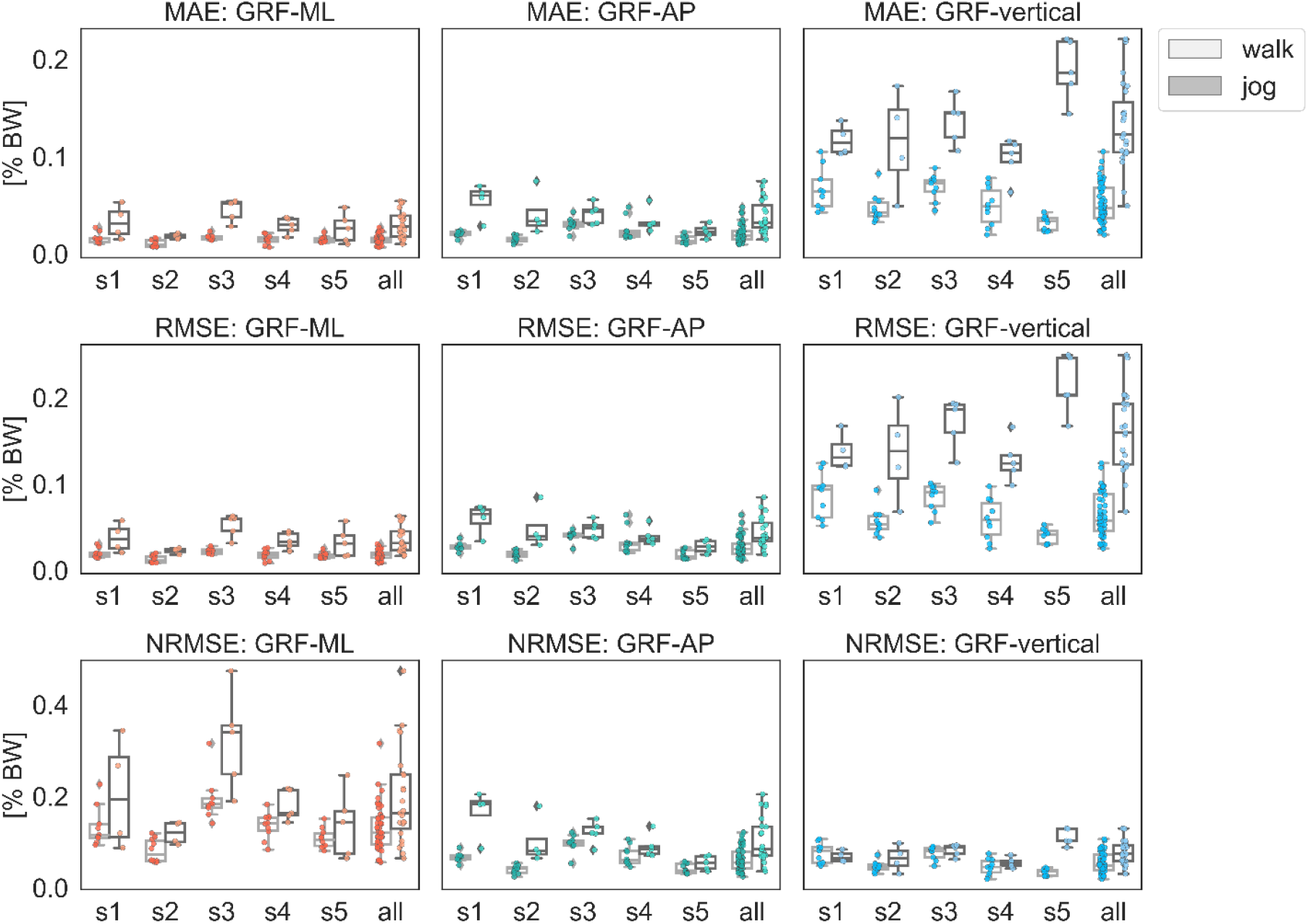
Box plots of the mean absolute error (MAE), root mean squared error (RMSE), and normalized root mean squared error (NRMSE) values of the ground reaction force prediction during walking and jogging. The accuracies are presented for each component of ground reaction force for each participant as well as all participants in the test set.

**Figure 4.**
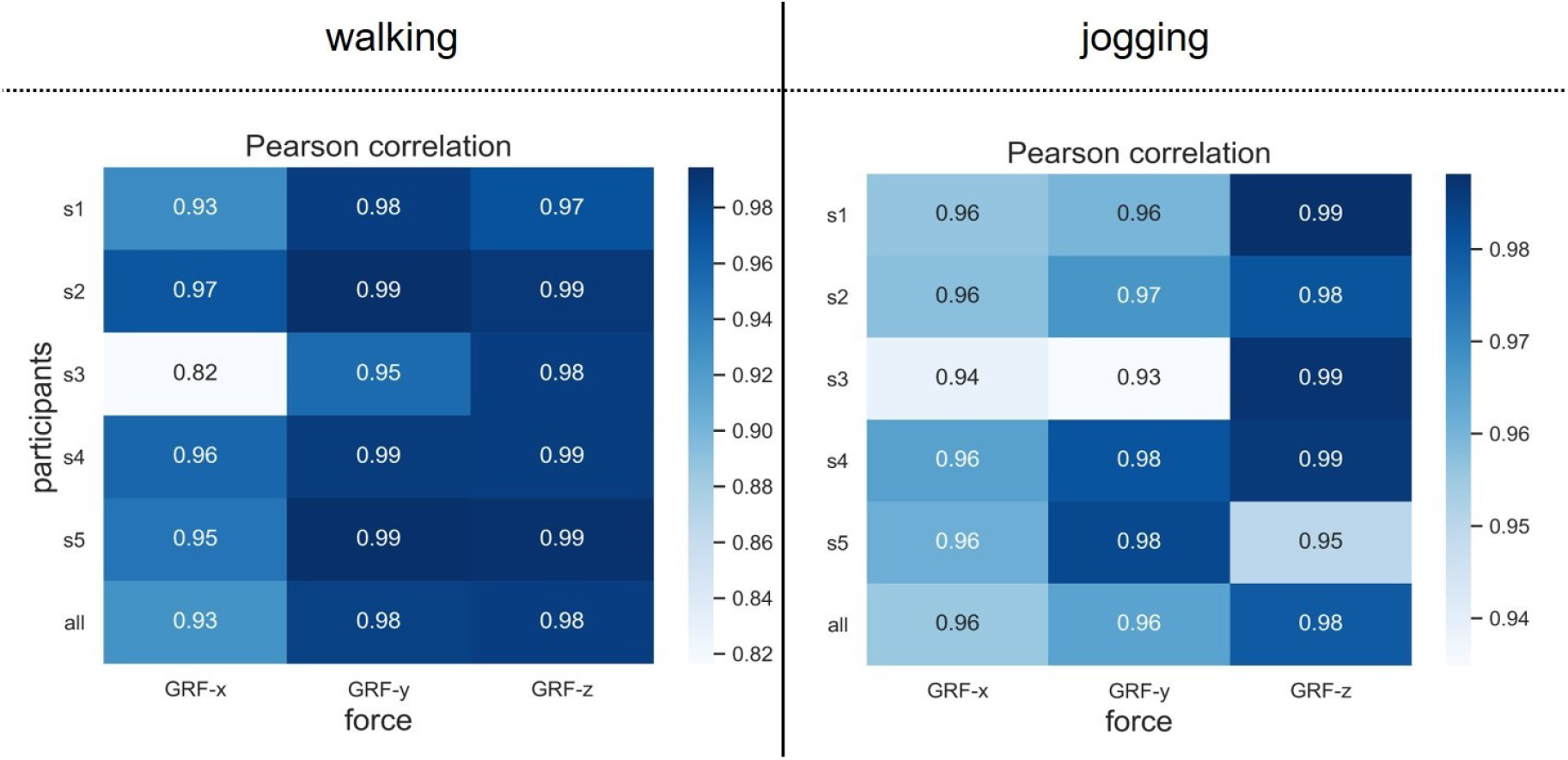
Correlation coefficient between the ground truth and predicted ground reaction force during walking and jogging. The correlations are presented for each component of ground reaction force for each participant as well as all participants in the test set.

In addition, the average errors during jogging were MAE= 13.25± 4.25% of body weight and NRMSE=7.72 ± 2.51% for vertical GRF, MAE= 3.82± 1.63% of body weight and NRMSE= 10.42± 4.75% for AP GRF, and MAE= 3.08± 1.39% of body weight and NRMSE= 19.57± 10.21% for ML GRF. The average correlation coefficient during jogging was 0.980± 0.019 for vertical force, 0.964± 0.030 for AP force, and 0.955± 0.021 for ML force. Finally, the performance evaluation metrics were summarized in Table 2 and compared with similar literature.

**Table 2.** Summary of the performance of different techniques to predict ground reaction force.

| Study | Model | Methods | GRF-x (ML)<br>Mean (SD) | GRF-y (AP)<br>Mean (SD) | GRF-z (V)<br>Mean (SD) |
| --- | --- | --- | --- | --- | --- |
| **NRMSE (%)** |  |  |  |  |  |
| This study | LSTM | Walking | 12.96 (4.98) | 6.26 (2.52) | 5.50 (2.25) |
| This study | LSTM | Jogging | 19.57 (10.21) | 10.42 (4.75) | 7.72 (2.51) |
| Rouhani et al.,<br>2010 | Linear | 11 participants (6 healthy, 5 ankle OA/<br>arthroplasty) | 21.58 (14.96) | 17.32 (9.06) | 15.51 (7.01) |
|  | MLP | 171 trials | 22.62 (13.32) | 16.37 (8.44) | 11.36 (4.74) |
|  | LLNF | 6-m walkway/ normal walking speed<br>Sandals | 21.58 (14.96) | 17.32 (9.06) | 15.51 (7.01) |
|  |  | Pressure insole (Pedar, Novel) |  |  |  |
| Sim et al, 2014 | WNN | 10 healthy participants<br>10 walking trials per subject<br>Participants' walking shoes<br>Pressure insole (Pedar, Novel) | 15.027 | 12.920 | 12.957 |
| **RMSE** |  |  |  |  |  |
| This study | LSTM | Walking | 1.81 (0.54) [% BW] | 2.75 (1.13) [% BW] | 6.45 (2.52) [% BW] |
| This study | LSTM | Jogging | 3.65 (1.50) [% BW] | 4.39 (1.66) [% BW] | 15.91 (4.48) [% BW] |
| Wei et al 2019 | Multi-stage<br>regression<br>(HS,<br>midstance,<br>TO) | 6 healthy participants<br>4.88 adjustable ramp<br>10 walking trials per subject<br>Merrell Trail Glove minimalist shoes<br>Pressure insole (Pedar, Novel) | 21 [% peak value] | 10 [% peak value] | 3 [% peak value] |
| Fong et al, 2008 | Stepwise<br>linear<br>regression | 10 healthy participants<br>10-m walkway<br>10 walking trials per subject<br>Standard sport shoes<br>Novel Pedar pressure insoles | 28 [% peak value] | 12 [% peak value] | 5 [% peak value] |

|  |  |  | Correlation coefficient |  |  |
| --- | --- | --- | --- | --- | --- |
| This study | LSTM | Walking | 0.926 (0.062) | 0.983 (0.018) | 0.985 (0.010) |
| This study | LSTM | Jogging | 0.955 (0.021) | 0.964 (0.030) | 0.980 (0.019) |
| Rouhani et al.<br>2010 | Linear |  | 0.764(0.259) | 0.906 (.072) | 0.967 (0.051) |
|  | MLP |  | 0.780 (0.172) | 0.908 (0.095) | 0.957 (0.046) |
|  | LLNF |  | 0.764 (0.259) | 0.906 (.072) | 0.967 (0.051) |
| Fong et al, 2008 | Stepwise<br>linear<br>regression |  | 0.719 | 0.928 | 0.989 |
| Sim et al, 2014 | WNN |  | 0.845 | 0.969 | 0.976 |
| Savelberg et al,<br>1999 | Feedforward<br>neural<br>network | 4 participants<br>Micro-EMED Novel pressure insole<br>Individuals' shoes<br>Walking | N/A | 0.621- 0.963 | N/A |

ML: mediolateral; AP: anteroposterior; V: vertical; SD: standard deviation; NRMSE: normalized root mean square error; RMSE: root mean square error
Neural network models: LSTM: long short-term memory; MLP: multilayer perceptron; LLNF: locally linear neuro-fuzzy; WNN: wavelet neural network

## 4. Discussion

The results of this study suggest that the LSTM model can be used to predict the 3D GRF from FScan plantar pressure insoles. The prediction results were validated for both walking and jogging, and the model performance was evaluated with four metrics: MAE, RMSE, NRMSE, and correlation coefficient. In general, the results showed high level of accuracy for 3D components of GRF during walking and jogging. However, the predicted shear forces were in better agreement with force plate data for walking compared to jogging, especially for AP forces.

The lowest NRMSE occurred for the vertical GRF, while the largest error was observed in ML GRF for both walking and jogging. The lower error being in vertical GRF compared to other components is in agreement with previous research [15, 17], and might be due to the lower variability of vertical GRF within and between participants compared to other GRF components [21]. During walking, our study showed lower values of NRMSE for predicting 3D GRF using an LSTM model compared to previous studies using other approaches such as multilayer perceptron networks [15, 16], linear neuro-fuzzy algorithms [15], and wavelet neural networks [17]. To our knowledge, previous studies did not predict GRF during jogging. The results of our study showed lower NRMSE for predictions of AP and vertical forces during jogging compared to previous research during walking. However, the NRMSE reported by Sim *et al*. [17] for predicting ML force during walking was lower than the corresponding component during jogging in this study. The correlation between the predicted and actual GRF was high for all three components, with highest correlation for vertical force and lowest correlation for ML force. Compared to previous studies, our results showed higher correlation for ML force during both walking and jogging. Regarding AP force, a slightly higher correlation coefficient (0.969) was achieved by Sim *et al*. [17] during walking compared to this study during jogging (0.964). Considering vertical force, our predictions were corelated to the actual forces with the average coefficients of 0.985 during walking and 0.980 during jogging, while Fong *et al*. reached a correlation coefficient of 0.989 during walking. The lower performance of the model during jogging compared to walking might originate from higher gait speed which could affect the variability of the 3D GRF [21]. In addition, the top surface of FScan pressure insoles is a thin plastic film (mylar sheets) which is flexible in one direction but does not conform well to 3D foot and shoe sole curvature [4, 22], is susceptible to sensor slipping [23], and may crinkle. These factors might have caused artefacts in the pressure data, especially during jogging [22].

An important achievement of this study was using the FScan pressure system with resistive sensors to predict forces without applying additional calibrations on pressure data. Based on the manufacturer’s guideline, additional calibrations have been suggested before recording data for each participant in addition to manufacturer’s calibration. Previous research has reported that the accuracy of FScan pressure system is highly dependent on these calibration and preconditioning sessions as they can reduce the effect of drift, creep, and hysteresis [9-11]. However, recording these further calibrations can be time consuming, and several fails in applying further calibrations might occur especially when the sensors have started to degrade. This complex process of calibration can be challenging or infeasible to perform in clinics and might consequently lead to wrong diagnosis [24]. Therefore, developing a model to use the pressure data without secondary calibrations and accurately estimate force can overcome such disadvantages and save time and computation costs. On the other hand, the accuracy of resistive sensors such as FScan system has been stated to be lower than the capacitive sensors such as the Pedar pressure system [4]. To the best of our knowledge, all the available literature that used commercial pressure sensors to predict GRF from pressure data have used Novel products including Pedar pressure insoles or Emed pressure mats. This is the first study to use FScan system with resistive sensors to demonstrate the possibility of accurate prediction of forces using these systems despite their limitations. It was observed that accurate predictions were feasible for all GRF components by implementing a robust deep learning model, with the added advantage that insoles with resistive sensors are much cost effective compared to the ones with capacitive sensors.

There are some potential limitations to this study. The findings of this study were reported for a group of healthy young participants, and the method has not been validated for different foot types, pathologies, or surface conditions. Future studies might generalize this model for different conditions such that a reliable prediction of GRF components from these wearable pressure insoles can be guaranteed. A further limitation is that the shoes properties were not the same between participants. This might have contributed to the variability between individuals and lower prediction accuracy during jogging. Future studies are recommended to use standard shoes during data collection. In addition, considering the force plates as gold standard to estimate the GRF from plantar pressure data might have not been ideal. While the foot is in direct contact with pressure insoles, the shoe sole is placed between foot and force plate. Therefore, the shoe material property might impact the GRF estimations by force plates.

The LSTM model developed in this study showed a good level of accuracy for predicting 3D GRF and has the potential to be integrated within the plantar pressure systems to enable real-time reflection of forces. The real-time reflection of the complete GRF with wearable plantar pressure insoles are more cost effective compared to force plates. They can also be used out of the laboratory to reflect the real-life forces during daily dynamic activities for both clinical and sport applications.

## Acknowledgement

This manuscript is a part of project which was financially supported by Mitacs. Funding agencies played no role in the design of the study, analysis, or writing of this manuscript.

