## Supplementary files for "Predicting vertical and shear ground reaction forces during walking and jogging using wearable plantar pressure insoles"

Summary:

Supplementary tables and figures that addressed in the body of the manuscript are presented here.

Supplementary Table 1: Summary of the accuracy for predicting ground reaction force during walking and jogging

| | | MAE [% BW] | RMSE [% BW] | NRMSE [%] | CC |
|---|---|---|---|---|---|
| walk | GRF-x (M/L) | 1.48 ± 0.45 | 1.81 ± 0.54 | 12.96 ± 4.98 | 0.926 ± 0.062 |
| | GRF-y (A/P) | 2.10 ± 0.86 | 2.75 ± 1.13 | 6.26 ± 2.52 | 0.983 ± 0.018 |
| | GRF-z (V) | 5.22 ± 2.05 | 6.45 ± 2.52 | 5.50 ± 2.25 | 0.985 ± 0.010 |
| Jog | GRF-x (M/L) | 3.08 ± 1.39 | 3.65 ± 1.50 | 19.57 ± 10.21 | 0.955 ± 0.021 |
| | GRF-y (A/P) | 3.82 ± 1.63 | 4.39 ± 1.66 | 10.42 ± 4.75 | 0.964 ± 0.030 |
| | GRF-z (V) | 13.25± 4.25 | 15.91 ± 4.48 | 7.72 ± 2.51 | 0.980 ± 0.019 |

MAE: mean absolute error; RMSE: Root mean squared error; NRMSE: normalized Root mean squared error; CC: correlation coefficient

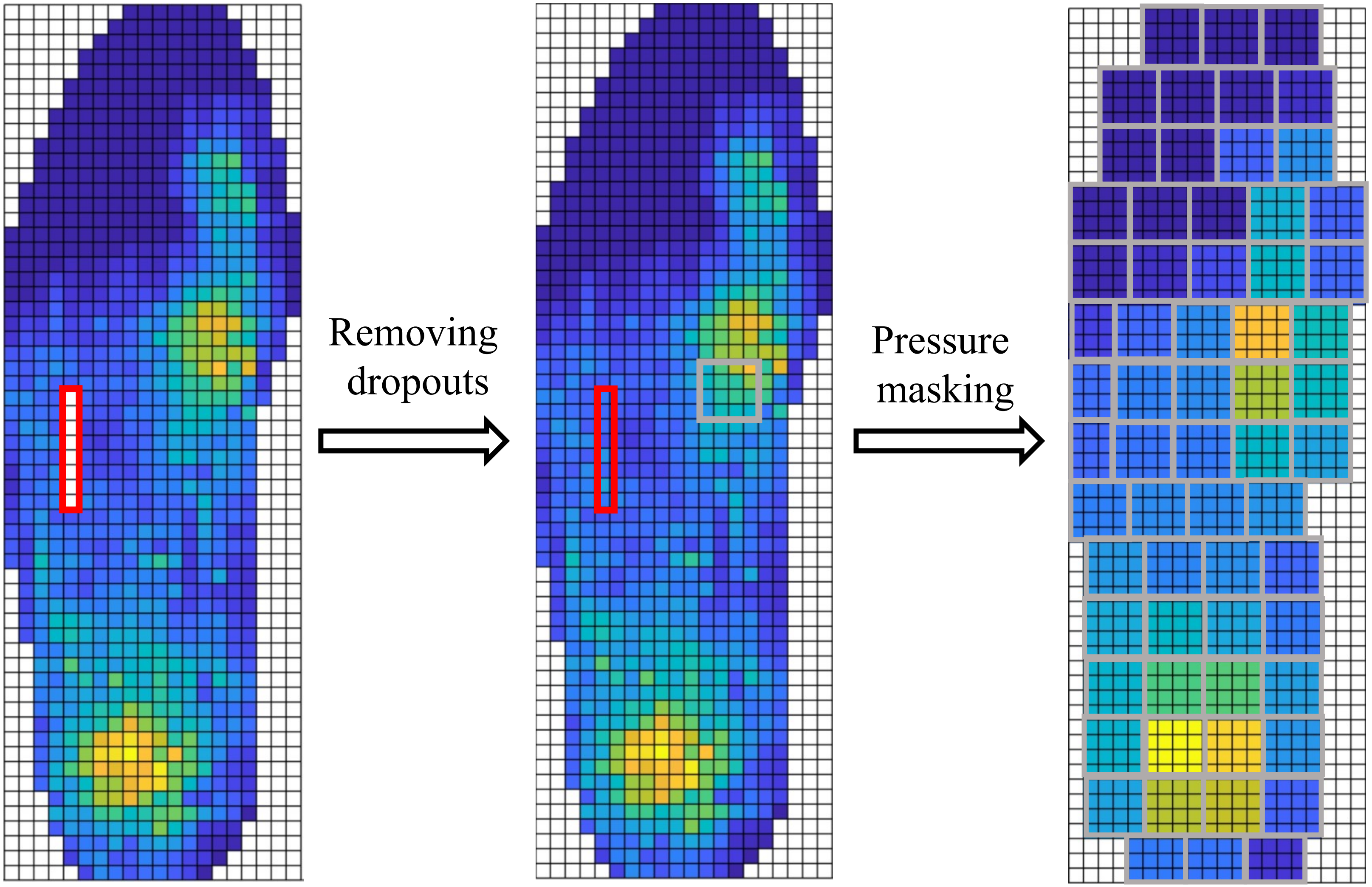


Supplementary Figure 1: The steps of masking of pressure data to prepare them for the input of neural network

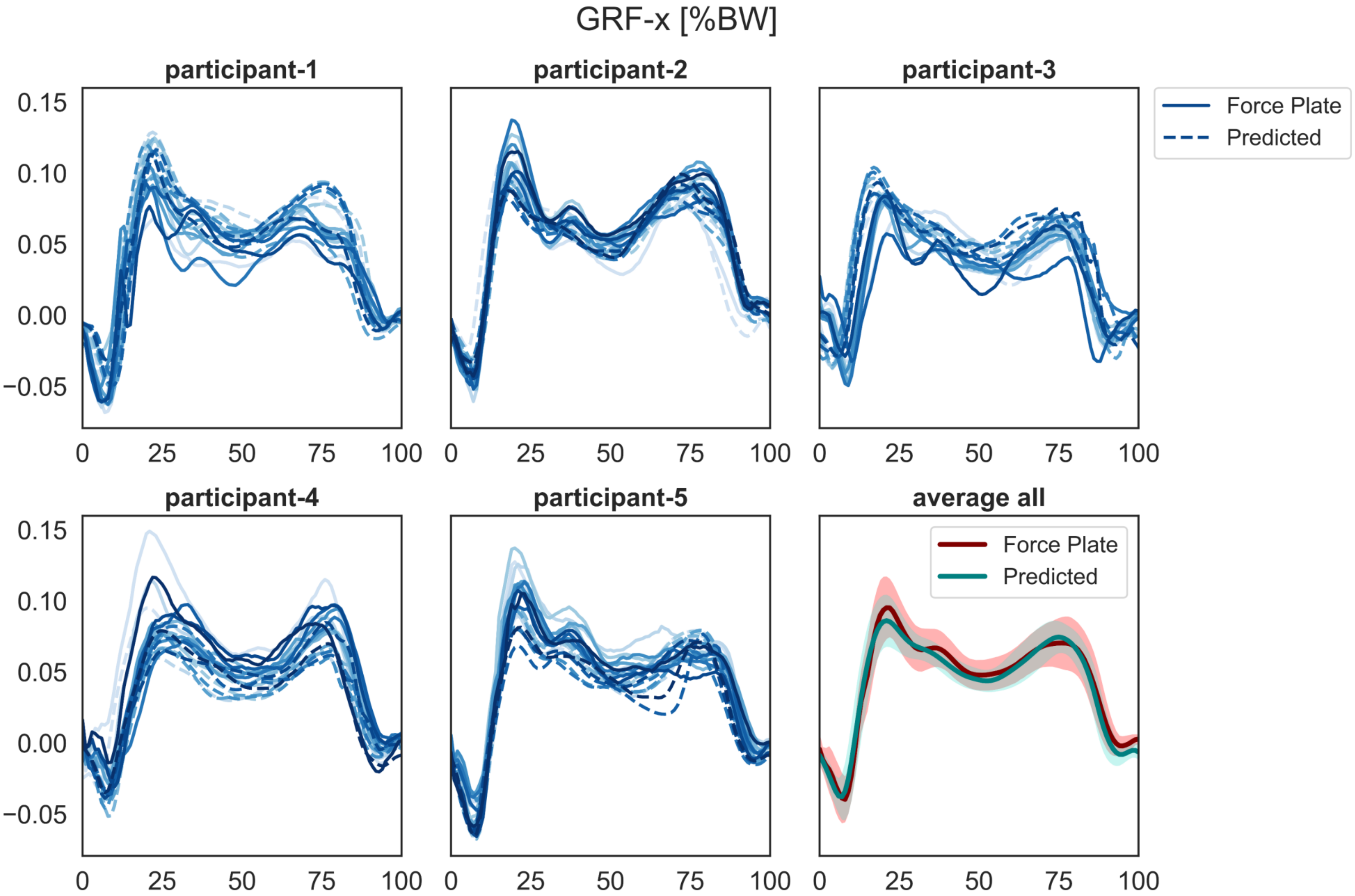


Supplementary Figure 2: The force plate versus predicted GRF-x (mediolateral component) for each participant in test set during walking

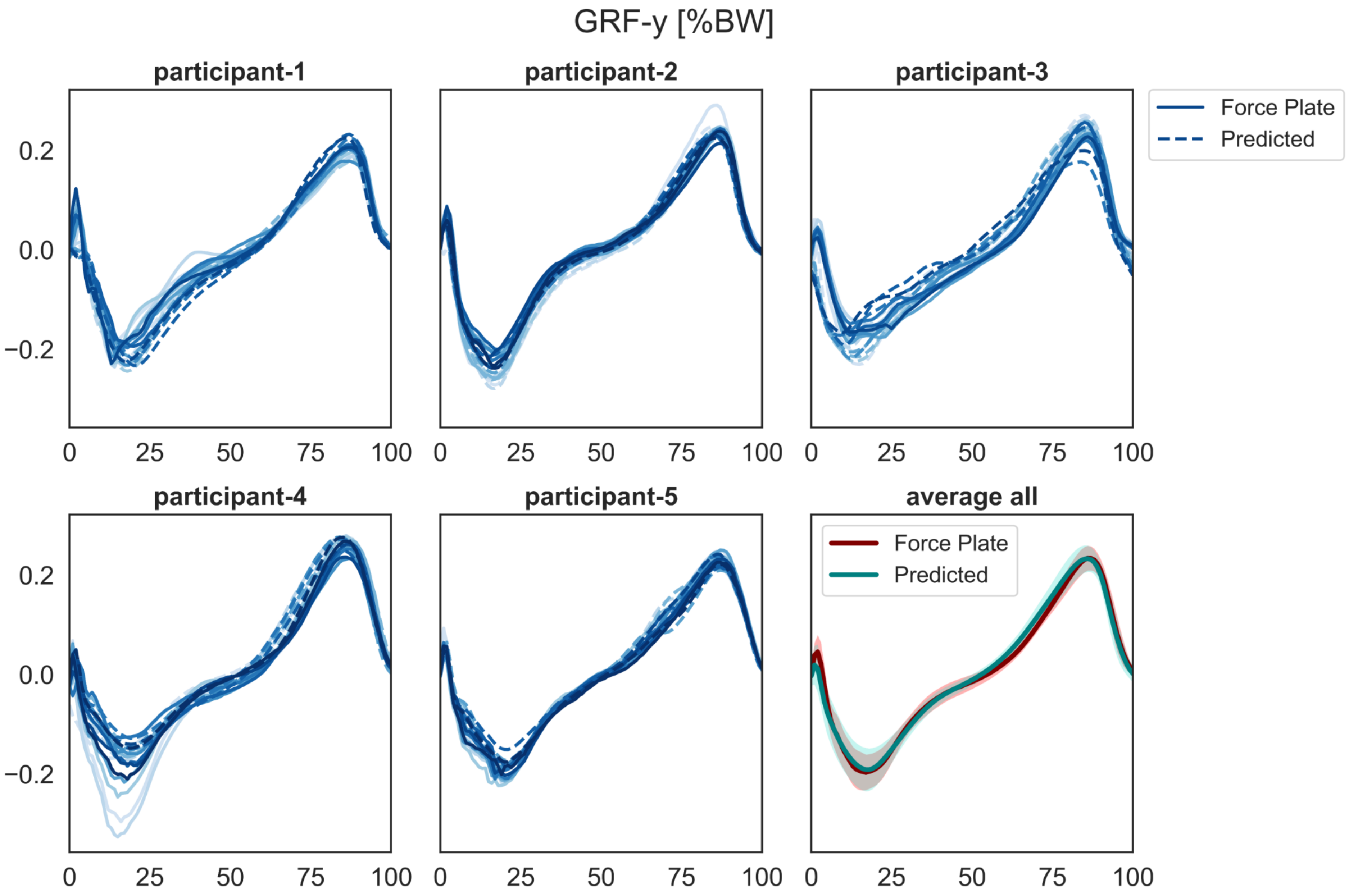


Supplementary Figure 3: The force plate versus predicted GRF-y (anteroposterior component) for each participant in test set during walking

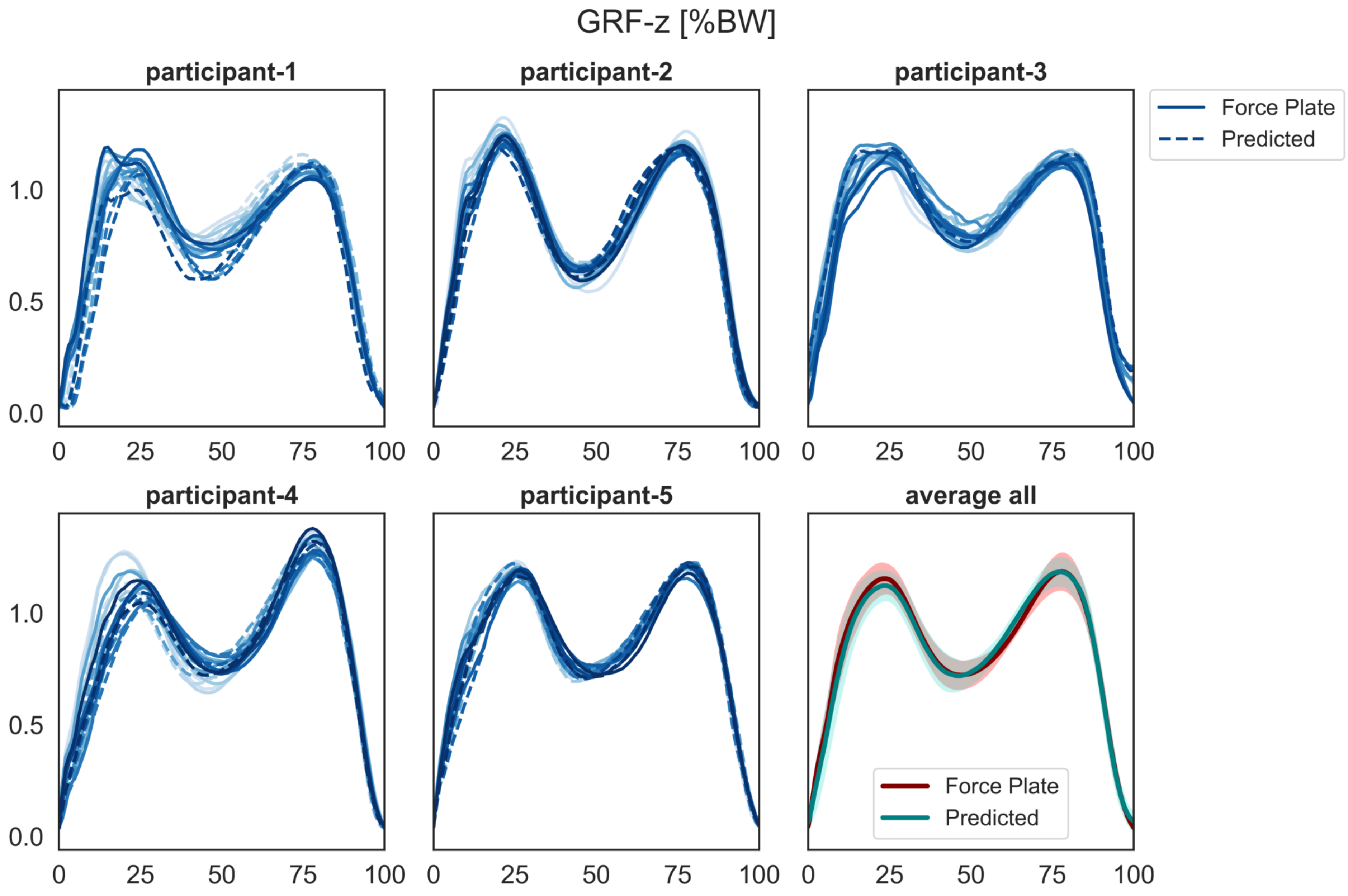

Supplementary Figure 4: The force plate versus predicted GRF-z (vertical component) for each participant in test set during walking

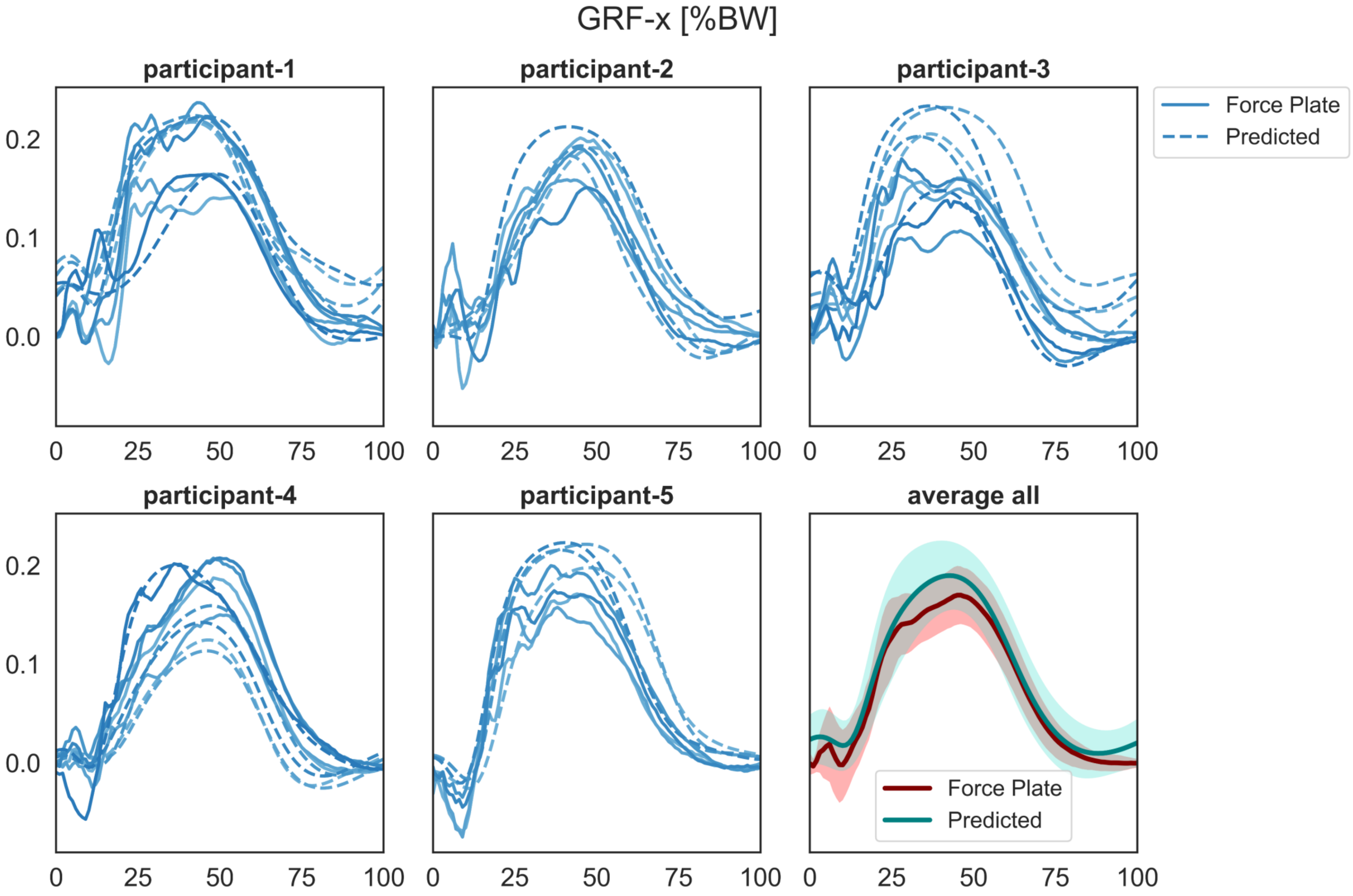


Supplementary Figure 5: The force plate versus predicted GRF-x (mediolateral component) for each participant in test set during jogging

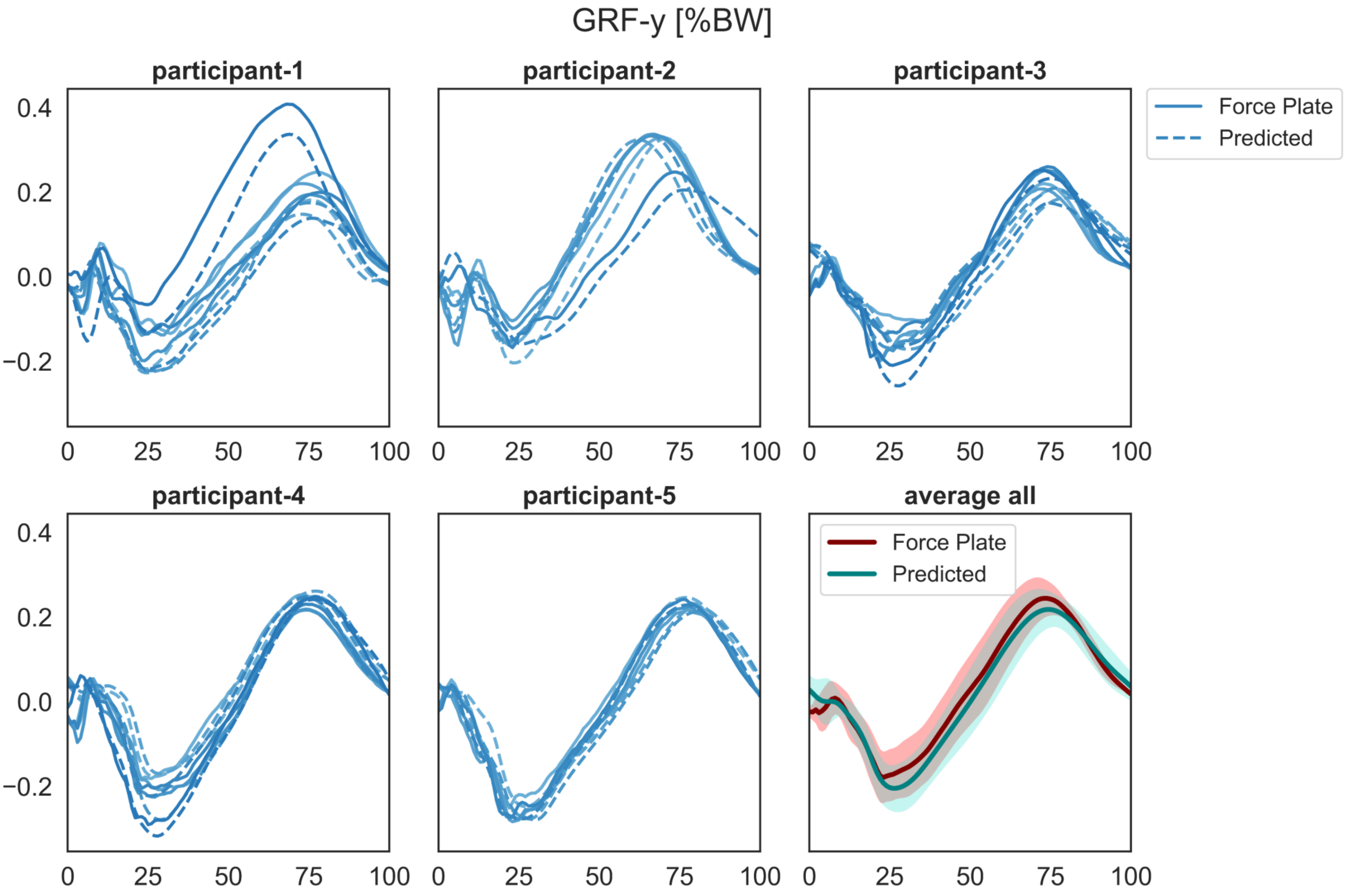


Supplementary Figure 6: The force plate versus predicted GRF-y (anteroposterior component) for each participant in test set during jogging

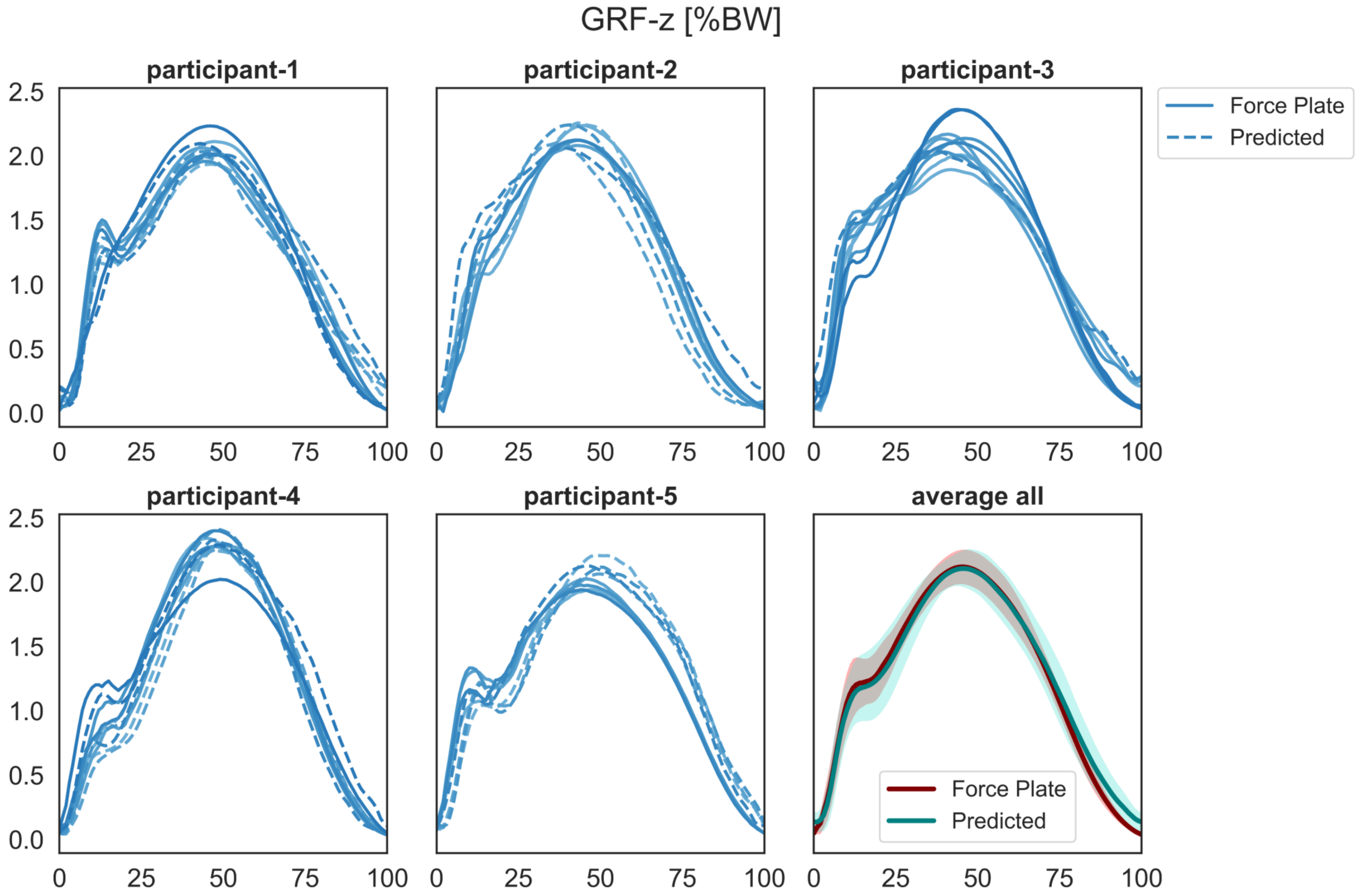


Supplementary Figure 7: The force plate versus predicted GRF-z (vertical component) for each participant in test set during jogging
